# A user-friendly machine-learning program to quantify stomatal features from fluorescence images

**DOI:** 10.1101/2025.11.20.689597

**Authors:** Gabriel J Angres, Alexander Gillert, Andrew Muroyama

## Abstract

In nearly all plants, pores on the leaf surface called stomata are essential for photosynthesis and gas exchange. The shape and distribution of stomata on the leaf varies widely between plants and is directly connected to photosynthetic efficiency. However, our understanding of the factors, both genetic and environmental, that exert subtle but significant effects on stomatal morphology is limited by the time required to manually annotate stomata in large imaging datasets. Here, we present a lightweight and efficient tool, QuickSpotter, for semi-automated stomatal annotation from fluorescence images. First, we establish QuickSpotter’s ability to automatically and accurately annotate mature stomata across developmental time. We also introduce an optional, speedy proofreading utility, StomEdit, that allows the researcher to quickly validate and correct machine-generated annotations. We use QuickSpotter and StomEdit to quantify how stomatal morphology evolves at the population level during cotyledon development and demonstrate how the programs can be used to extract subtle differences in stomatal development following pharmacological treatments. Finally, we describe PairCaller, a pair-calling classifier that accompanies QuickSpotter and can be used to identify stomatal clusters, a physiologically relevant and widely studied developmental phenotype. Taken together, our suite of programs facilitates quantitative analyses of stomatal development at scale, enabling high-throughput analyses of leaf phenotypes under varied conditions.

## Introduction

Stomata are essential pores on the surfaces of plant leaves that regulate photosynthesis and transpiration. While the basic stomatal structure has been conserved for hundreds of millions of years (Edwards *et al*., 1998), stomata display significant morphological variability between different species, ecotypes, environmental conditions (Hetherington & Woodward, 2003; Rudall *et al*., 2017; Bertolino *et al*., 2019; Pan *et al*., 2024). As such, stomatal size and distribution in the epidermis are directly coupled to photosynthetic efficiency, as evidenced by recent work demonstrating that eudicots and monocots with altered stomatal density or clustering exhibit marked differences in photosynthetic capacity (Schlüter *et al*., 2003; Dow *et al*., 2014; Lehmann & Or, 2015; Harrison *et al*., 2020; Xiong *et al*., 2022; Haworth *et al*., 2023). Because they directly impact water-use efficiency, there are ongoing efforts to engineer stomatal characteristics to improve crop tolerance to environmental stress (Endo & Torii, 2019; Bouvier & Kelly, 2025). Therefore, there remains significant interest in understanding both the genetically encoded and environmentally responsive pathways that control stomatal development.

Many studies continue to use image-based stomatal phenotyping, including measures of size, density, and pairing, to identify mutations, environmental conditions or abiotic and biotic stresses that impact stomatal development. There remain both biological and technical hurdles to efficient workflows capable of handling this type of data. In species such as *Arabidopsis*, where the majority of this research continues to be conducted, stomatal distribution and orientation are unique and variable in each leaf, making it impossible to use global coordinates to systematically predict stomatal positions. Therefore, manual annotation remains the standard in the field, but even with dedicated, open-source image-analysis tools like Fiji (Schindelin *et al*., 2012), this process requires a substantial time investment. While simply counting stomata in a small field of view can be accomplished in a few minutes per image, manually tracing stomatal areas for precise morphological analyses exponentially increases the time required, creating a significant bottleneck. The issue is further compounded when researchers are interested in quantifying all stomata within a given leaf, for example when asking questions related to stomatal heterogeneity across the leaf surface (Weyers *et al*., 1997) or the effects of stomatal distribution on photosynthesis (Harrison *et al*., 2020). Therefore, there remains a pressing need to develop automated or semi-automated means of identifying and quantifying stomata at scale.

Machine vision (MV) represents a promising solution to this bottleneck in image-based phenotyping. Since their inception, MV techniques have gained distinction as an approach to conducting accurate and high-throughput visual analysis of biological phenomena, and for good reason: at their best, machine-aided image analysis tools can perform at or above the level of human experts while operating in a fraction of the time and with minimal human oversight. In recent years, these tools have been used for applications in plant biology as varied as identifying diverse diseases and deficiencies in soybean leaves (Ghosal *et al*., 2018), connecting weather conditions with cotton pest activity (Xiao *et al*., 2019), and estimating melon harvest weight through unmanned aerial vehicle (UAV) imaging (Kalantar *et al*., 2020). By utilizing publicly available frameworks that permit the application of MV in nearly any imaging context, development of a MV pipeline to perform high-throughput stomatal characterization would greatly expedite ongoing and future work on stomatal development and physiology.

Several groups have previously reported programs that facilitate imaging-based stomatal phenotyping. Some of these programs use brightfield images of optically cleared samples as their sole input (Vialet-Chabrand & Brendel, 2014; Jayakody *et al*., 2021; Wang *et al*., 2024). These are particularly useful for applications in field studies and non-model plant species, from which fixed and cleared samples can be readily prepared. However, these types of programs are limited in their ability to analyze developmentally resolved data from the same sample. Other programs take brightfield images or scanning electron micrographs as an input (Fetter *et al*., 2019; Wolny *et al*., 2020; Li *et al*., 2022), but are more suited to stomatal counting than the generation of high-accuracy masks. Finally, still others, such as PaCeQuant (Möller *et al*., 2017) and MorphoGraphX (Barbier de Reuille *et al*., 2015; Strauss *et al*., 2022) are particularly useful for analyzing pavement cells but show a more limited ability to accurately segment stomata. Part of the challenge associated with developing a program capable of identifying stomata is that the cellular makeup of the leaf epidermis changes dramatically during leaf development. Early stomatal lineage cells are highly abundant in early stages, when pavement cells are also relatively small, while the epidermis is composed almost entirely of large, lobed pavement cells and stomata at later stages. Therefore, one of our goals was to develop a program capable of robustly segmenting stomata regardless of developmental stage.

In this paper, we report the development of QuickSpotter, a semantic segmentation algorithm for stomatal annotation from confocal images that can 1) accurately segment stomata for morphological analyses, 2) enable both fully automated and semi-automated segmentation, 3) empower program re-training for more specialized applications and 4) run efficiently on a typical lab computer without the need for expensive setups or high-performance computer cluster access.

Here, our goal was to create an accurate and user-friendly MV model from pretrained weights, both to segment stomata and to classify whether a given annotation feature belonged to a single stoma or a cluster of two or more adjacent stomata. After validation, we applied the model, along with a custom-built graphical user interface (GUI) for proofreading machine-generated annotations, to comprehensively benchmark stomatal number and morphology during Col-0 cotyledon development. Furthermore, we use our tool to perform phenotypic characterization of stomata following pharmacological treatments, highlighting its potential for enabling biological discovery in large image datasets.

## Materials and Methods

### Plant lines and growth conditions

ML1p::mCherry-RCI2A (Roeder *et al*., 2010) and *basl-2* 35Sp::PIP2A-RFP (Rowe *et al*., 2019) lines were previously reported. Seeds were sterilized in 20% bleach containing 0.01% Triton-X for 10 minutes then washed three times with sterilized ddH2O before plating on ½ MS + 0.5% sucrose plates. Plated seeds were stratified in the dark at 4°C for at least 2 days before being grown in a Percival growth chamber set to 22°C with a 16hr/8hr light/dark cycle. For AZD-8055 treatments, seedlings were stratified and grown on ½ MS + 0.5% sucrose plates containing the indicated concentrations of AZD-8055 (MedChemExpress).

### Acquiring Images

All images were acquired on a Leica Stellaris 5 scanning confocal microscope equipped with HyD detectors. All images were acquired using a 20x/0.75NA objective.

### Image Processing

Acquired LIF image files were converted to single-layer TIFs using Fiji’s Z-Project>>Max function on all layers. Initial annotation masks, except where otherwise noted, were produced manually using native drawing functionality in Paint.NET (dotPDN LLC, 2025). All annotation overlays denoting stomatal positions (hereafter referred to as “masks”) used for training and validation were formatted as solid black and white (#000000 and #FFFFFF color values) PNG files.

### Python Libraries and Framework

The machine-learning algorithm of choice, implemented through PyTorch (Paszke *et al*., 2019), was a U-Net architecture (Ronneberger *et al*., 2015), modified with batch normalization layers (Ioffe & Szegedy, 2015) and a MobileNetV3-Large backbone (Howard *et al*., 2019) in place of U-Net’s default starting configuration. This framework configuration (including architecture and weights) was optimized for its performance on biological image segmentation tasks while simultaneously being lightweight and efficient (Ronneberger *et al*., 2015). The counterpart proofreading GUI was developed with the TKinter package. Accompanying image processing and display was implemented with the Scikit-Image (SKImage) and PILLOW libraries.

### Training and Annotation Cycles

Training data preparation was initialized with pairs of “base” and “annotation” files corresponding to the raw LSCM images and their annotated black-and-white masks, respectively. The masks were drawn such that overlaying the mask onto the base image would exactly occlude the stomata present on the cotyledon with white blotches.

First, each image pair was broken down into pairs of corresponding 128×128 pixel squares, tiled with overlap over both images. To increase the number of tiles generated from each image without creating redundant training data, the windows overlapped each other by a 1/8^th^ of their width (16 pixels), increasing the tiles generated by ∼30%. This ensured that any one stoma would always be present in at least one image without being clipped by the window border, improving the depth of the training data.

Additional data augmentation steps were used to expand the training set. All pairs of images underwent a quarter turn applied 1-4 times and had an equal chance to be flipped on the vertical axis or left unmodified. From there, each image had a one in three (1/3) chance to undergo a Gaussian blur transformation with a random radius and intensity (kernel from [3,5,7], sigma from [0.0-3.0]). While blurs of this intensity do not necessarily occur routinely *in situ*, these transformations were included to improve the model’s resilience to variation in image quality.

Hence, each 128×128 section could appear as one of 8 different variants before blurring was applied. However, rather than generating every variant of every image for each training cycle, one variant of each individual segment was generated at random for each epoch, changing the input dataset between epochs. Theoretically, this choice makes training cycle performance non-deterministic or probabilistic (variable), reducing the risk of overfitting but with potential impacts on generalized performance.

In addition, each model also required a selection of hyperparameters, including the algorithm learning, the pixel dimensions of the examined window, and the overlap proportion between each sample window. These settings were iteratively tuned and evaluated to create an optimal balance between training time and fit quality.

A higher learning rate increases the likelihood of convergence but has a greater probability of overfitting, whereas a lower learning rate would be more conservative but likely fall short of an optimal convergence. Increasing the window size would provide more data for each training step but increase training time and increase the chance of overfitting. Ultimately, a 128×128 window size was chosen to ensure that at least one stoma and no more than three or four would ever be in the same viewing window at once, reducing training time and the possibility of overfitting. Finally, the degree of overlap increased the amount of input data and training time but also allowed a given region of the image to be represented multiple times in different contexts.

The model was trained to identify two classes in its segmentation: whether the pixel belonged to a stoma (“1”, white, “true”) or not (“0”, black, “false”). Over each pixel in a given region of the image, the algorithm assigned a value on the range (-inf, inf), with values farther from 0 becoming exponentially less common, and with no values over magnitude 12 observed in practice. A sigmoidal function converted these values to a range of 0-1, which were subsequently normalized to the range 0-255 to be displayed in grayscale. As judgements from multiple windows made multiple contributions to pixels, these judgements are combined to produce a composite grayscale whole-image prediction. Hence, the primary output of the model is a grayscale overlay with the same image size and resolution as the base image. Pixel values in the overlay range between 0 and 255, corresponding to the confidence that the algorithm has that a particular pixel covers an area within a stoma. A value of 0 (“black”) or 255 (“white”) indicates 0% and 100% confidence, respectively, with all intervening values corresponding to a sliding confidence scale. While this mask can be used as-is for visualization purposes, the overlay may be processed to a binary image with pixel values of either 0 or 255 to enable speedy and accurate stomatal quantification, which will be discussed further below.

### Training/Validation Set Construction

#### Stomatal Pixel Annotator

To train the model, two types of input image were used: “whole” images consisting of whole-leaf images and a corresponding mask, and “segments,” 512 pixel x 512 pixel sections with a corresponding 512×512 mask. The training set used 142 segments from Col-0 cotyledons at 3, 4, 5, and 7dpg, as well as 2 full-size 5dpg *basl-2* cotyledons and 12 full-size wild-type cotyledons (eight 3dpg, two 4dpg, two 5dpg). The validation set used one whole 3dpg Col-0 cotyledon, two segments of 3dpg Col-0 cotyledons, three segments of 7dpg Col-0 cotyledons and one whole 5dpg *basl-2* cotyledon. Validation data was set aside from the given stages to intentionally test the model performance on extremes of stomatal development. The dataset makeup for the training and validation runs is listed in Supplemental Table 1.

#### PairCaller Classifier

To complement the automatic stomatal annotator, a separate classifier algorithm, based on the MobileNetV3 network (Howard *et al*., 2019) implemented in Pytorch, was trained to determine whether a given annotation corresponded to a single or clustered stomata. All images used for training and validation were 72×72 pixels, since this size was sufficient to capture most single and clustered stomata. The training data contained a total of 3436 images, with 2870 images containing single stomata and 566 images containing clusters. The validation set contained a total of 879 images, with 714 images containing single stomata and 165 images containing clustered stomata.

### Image Post-Processing

After each machine prediction, several post-processing steps were applied to repair defects in the image. First, any object larger than 3750 pixels was removed to avoid large artifacts. This threshold was chosen because that area exceeds the size of even the largest observed hydathode stomata or stomatal clusters in the dataset. After Otsu thresholding was applied to convert the masks to binary images, an area dilation/erosion was conducted to remove any small, branch-like projections from the masks. Finally, any object smaller than 250 pixels was removed, as stomata smaller than this were not found in our dataset, and all dark holes within the annotation were filled-in. This small object removal step was conducted last to address artifacts that could be formed during the dilation/erosion step.

### Statistical analysis

Statistical analysis was performed in RStudio (Posit team, 2025). The IoU metrics were calculated in Python. Kruskal-Wallis tests were used to evaluate global differences between treatments. Dunn’s tests were used for post-hoc testing, with p-values adjusted by Bonferroni’s correction to reduce the family-wise error rate. Ridgeplots were generated in R, and stomatal density graphs were plotted using GraphPad Prism 10.6.1 (GraphPad Software, Boston, Massachusetts, USA, www.graphpad.com).

## Results

### Model Training

Initially, we attempted to build a model to identify all stomata, from the youngest guard cells that lack pores to fully mature stomata with well-defined pores, by annotating all instances of stomata in our training images. However, early training cycles showed that a segmentation model trained on these data struggled to consistently differentiate *bona fide* guard cell pairs from young pavement cells or early stomatal lineage cells that shared some morphological characteristics with immature stomata. Based on these preliminary trials, we modified our approach and decided to train our model exclusively on annotated stomata with visible pores. As pore formation commences within several hours of cytokinesis completion (Rui *et al*., 2019), we decided that this was a reasonable tradeoff to significantly improve model accuracy. Models developed to segment these pore-containing stomata (hereafter referred to collectively as “stomata”) improved prediction consistency and significantly reduced the rate of false positives (Figure 1).

**Figure 1.**
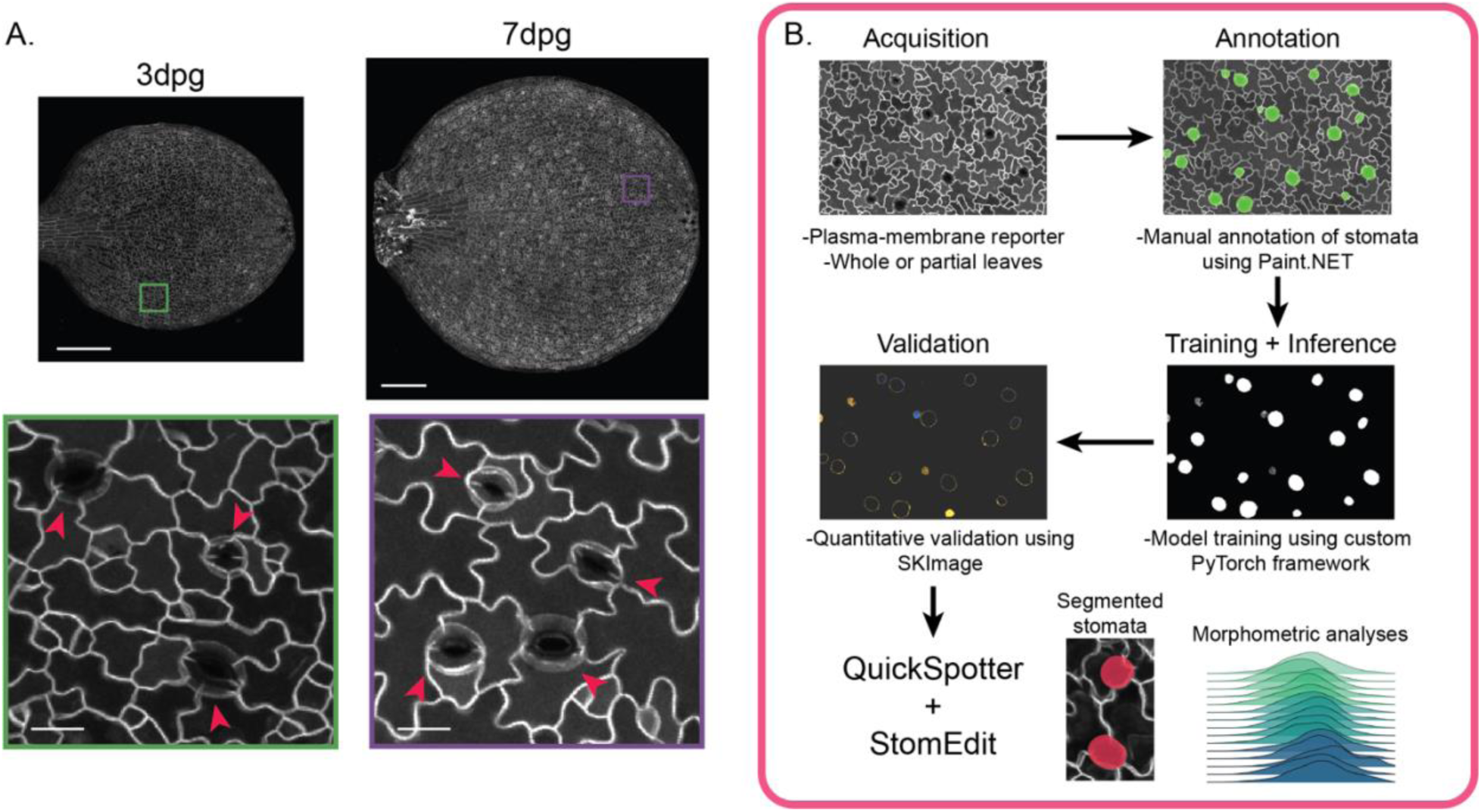
Development of an automatic stomatal annotation pipeline for morphometric analysis. A. Representative images of whole 3dpg and 7dpg Col-0 cotyledons. Mature stomata are marked with red arrows. Scale bars – 250μm (top) and 20μm (bottom). B. Training and configuration pipeline for QuickSpotter. CLSM images taken of a plasma membrane reporter were acquired and then manually annotated at the pixel level using Paint.NET, a free image-editing software. Annotation masks were subsequently paired with their base images and used as input for training/testing cycles to configure the MobileNetv3-backed U-NET neural net. After the final model was assessed for human-level quantitative and qualitative accuracy using Python’s SKImage, the final trained model, along with optional proofreading with StomEdit, can be used for mask generation and morphometric analyses.

We trained the model on 3,000+ manually traced stomata from greyscale images of 3, 4, 5 and 7 days post-germination (dpg) Col-0 cotyledons that expressed an epidermis-specific plasma membrane reporter, ML1p::mCherry-RCI2A. Images were curated from multiple imaging sessions with varied acquisition conditions to increase prediction robustness (Table 1). To accurately evaluate model performance, multiple training/validation runs were performed for each set of hyperparameters, such as the learning rate, using the same annotated training data (Figure S1). After several rounds of hyperparameter tuning focused on optimizing validation set convergence, we found that a window size of 128×128 pixels with 16-pixel overlap (roughly ⅛ of the window) with a learning rate of 0.001 provided sufficiently rapid convergence without overfitting.

**Table 1.** Samples used for QuickSpotter training and validation. Each cell corresponds to the number of stomata present in the training set from a segmented or whole image (rows), arranged by developmental age in dpg (columns) and whether they are present in the training or validation set (training/validation). Of note: single stoma and clusters from *basl-2* cotyledons were included in the training and validation data but are not included in this table for ease of presentation. See Table 4 for information regarding the *basl-2* contributions.

|  | 3dpg | 4dpg | 5dpg | 7dpg |
| --- | --- | --- | --- | --- |
| Segments | 579 / 25 | 226 / - | 250 / - | 431 / 34 |
| Whole Images | 642 / 112 | 442 / - | 784 / - | - / - |

### Benchmarking stomatal identification

The algorithm was evaluated on its performance on two key tasks: feature clarity and per-pixel accuracy. Feature clarity is a measure of the algorithm’s qualitative prediction capacity, in this case its ability to identify and create masks for stomata on the level of a human expert. Meanwhile, per-pixel accuracy is a quantitative metric that determines how many pixels in each stomatal annotation were mislabeled compared to the human-defined ground truth annotation. These two metrics, therefore, provide useful measures of how well the program can identify stomata in each image.

#### Feature Clarity

We first evaluated how well our program identified stomata in images that were set apart from the training dataset and, therefore, untouched during the training phase. When evaluating the model’s predictions, we classified each annotation as “Complete,” “Partial,” “False,” or “Miss,” defined as:

- **Complete** - an annotation whose convex boundary differs from the real stomatal outline by no more than three pixels at any point.
- **Partial** – 1) an annotation whose convex outline fails to capture the stomatal boundary and exceeds the boundary by three pixels or more at any point or 2) an annotation whose concave outline exceeds the stomatal boundary by three pixels or more.
- **False** – an annotation that does not correspond to a stoma.
- **Miss** – a stoma that is completely missed by the program.

For the purposes of assessing program accuracy, both Complete and Partial predictions were scored as positive identifications. Table 2 shows the results from the validation tests using a representative set of images, without any additional human correction. No “False” results were obtained in this data, but we noted that the image acquisition parameters did affect the rate of “False” results; issues such as under- or over-exposure, which generally affect image usability and quantification, influenced the rate of “False” calls, highlighting the importance of standardizing imaging conditions. Overall, the program successfully identified 92% of the stomata in 40 images (550 out of 586 stomata). The program missed between 0-3 stomata per image, with an average of 0.9 misses over all the analyzed images. Additionally, only two images produced a single False result, with an average of 0.05 False results over all images.

**Table 2.** Accuracy rates of QuickSpotter-identified stomata. The table shows Hit, Miss, and Partial counts for each of the 40 images used for QuickSpotter validation.

| Sample ID | Hit | Miss | Partial | Total | % Accurate |
| --- | --- | --- | --- | --- | --- |
| 5dpg_01 | 19 |  |  | 19 | 100% |
| 5dpg_02 | 12 | 1 |  | 13 | 92% |
| 5dpg_03 | 14 | 1 | 1 | 16 | 88% |
| 5dpg_04 | 17 | 2 |  | 19 | 89% |
| 5dpg_05 | 12 | 1 | 1 | 14 | 86% |
| 5dpg_06 | 15 |  |  | 15 | 100% |
| 5dpg_07 | 11 |  | 3 | 14 | 79% |
| 5dpg_08 | 12 | 2 | 2 | 16 | 75% |
| 5dpg_09 | 14 |  |  | 14 | 100% |
| 5dpg_10 | 17 |  | 1 | 18 | 94% |
| 5dpg_11 | 13 | 1 | 1 | 15 | 87% |
| 5dpg_12 | 13 | 1 | 1 | 15 | 87% |
| 5dpg_13 | 13 |  |  | 13 | 100% |
| 5dpg_14 | 11 |  | 4 | 15 | 73% |
| 5dpg_15 | 17 |  | 2 | 19 | 89% |
| 5dpg_16 | 16 |  |  | 16 | 100% |
| 4dpg_01 | 16 |  | 1 | 17 | 94% |
| 4dpg_02 | 11 | 3 | 1 | 15 | 73% |
| 4dpg_03 | 13 | 1 |  | 14 | 93% |
| 4dpg_04 | 14 | 2 |  | 16 | 88% |
| 4dpg_05 | 14 | 2 | 2 | 18 | 78% |
| 4dpg_06 | 17 |  | 2 | 19 | 89% |
| 4dpg_07 | 11 | 1 |  | 12 | 92% |
| 4dpg_08 | 14 |  | 3 | 17 | 82% |
| 4dpg_09 | 14 | 2 |  | 16 | 88% |
| 4dpg_10 | 11 | 3 | 1 | 15 | 73% |
| 4dpg_11 | 12 |  | 2 | 14 | 86% |
| 4dpg_12 | 15 | 3 | 1 | 19 | 79% |
| 3dpg_01 | 10 | 1 |  | 11 | 91% |
| 3dpg_02 | 10 |  |  | 10 | 100% |
| 3dpg_03 | 11 |  | 1 | 12 | 92% |
| 3dpg_04 | 9 | 1 | 4 | 14 | 64% |
| 3dpg_05 | 11 | 1 |  | 12 | 92% |
| 3dpg_06 | 9 | 1 | 2 | 12 | 75% |
| 3dpg_07 | 9 | 3 | 3 | 15 | 60% |
| 3dpg_08 | 7 |  | 1 | 8 | 88% |
| 3dpg_09 | 8 |  |  | 8 | 100% |
| 3dpg_10 | 8 | 2 | 3 | 13 | 62% |
| 3dpg_11 | 11 |  | 2 | 13 | 85% |
| 3dpg_12 | 13 | 1 | 1 | 15 | 87% |
| TOTAL | 504 | 36 | 46 | 586 | 86.01% |

#### Per-Pixel Accuracy

Next, we evaluated the per-pixel accuracy of the machine-generated masks by calculating the Intersection over Union (IoU) score, which is derived by dividing the pixel overlap between the ground-truth annotation and machine prediction by the combined total pixel area of the annotation and prediction. The IoU score can range from 0 to 1, indicating a complete mismatch or match, respectively. After calculating a contingency table for all masks in the ground truth data and the machine predictions and determining the degree of overlap between them, the metrics were aggregated into a single score to show the average image-wide accuracy (Table 3). To aid visual validation of the results, a function was programmed that overlaid the ground truth mask (as a black-and-blue image) with the machine prediction (as a black-and-orange image) to show the degree of accuracy. Using this scheme, all false negatives are blue, false positives are orange, true negatives are black, and true positives are the overlap between blue and orange and shown in pink (Figure 2 D,E). The average IoU score across the entire dataset, which included both tiled images of whole cotyledons and smaller fields of view within a single cotyledon, was 83.2% (range 61.1-95.2%). Excluding image segments that had stomata on the edges of the image, which are harder to predict owing to their partial nature, yielded a mean of 89.2% (range 80.6-95.2%). We noted that fluorescence intensity can vary across the cotyledon surface in some tiled images, as expected owing to subtle but meaningful differences in sample distance from the objective. Accordingly, accuracy on whole cotyledon images, which do not include any clipped stomata but have more heterogeneity in fluorescence intensity than single fields of view, was 87.8% (range 80.6 to 90.3%).

**Figure 2.**
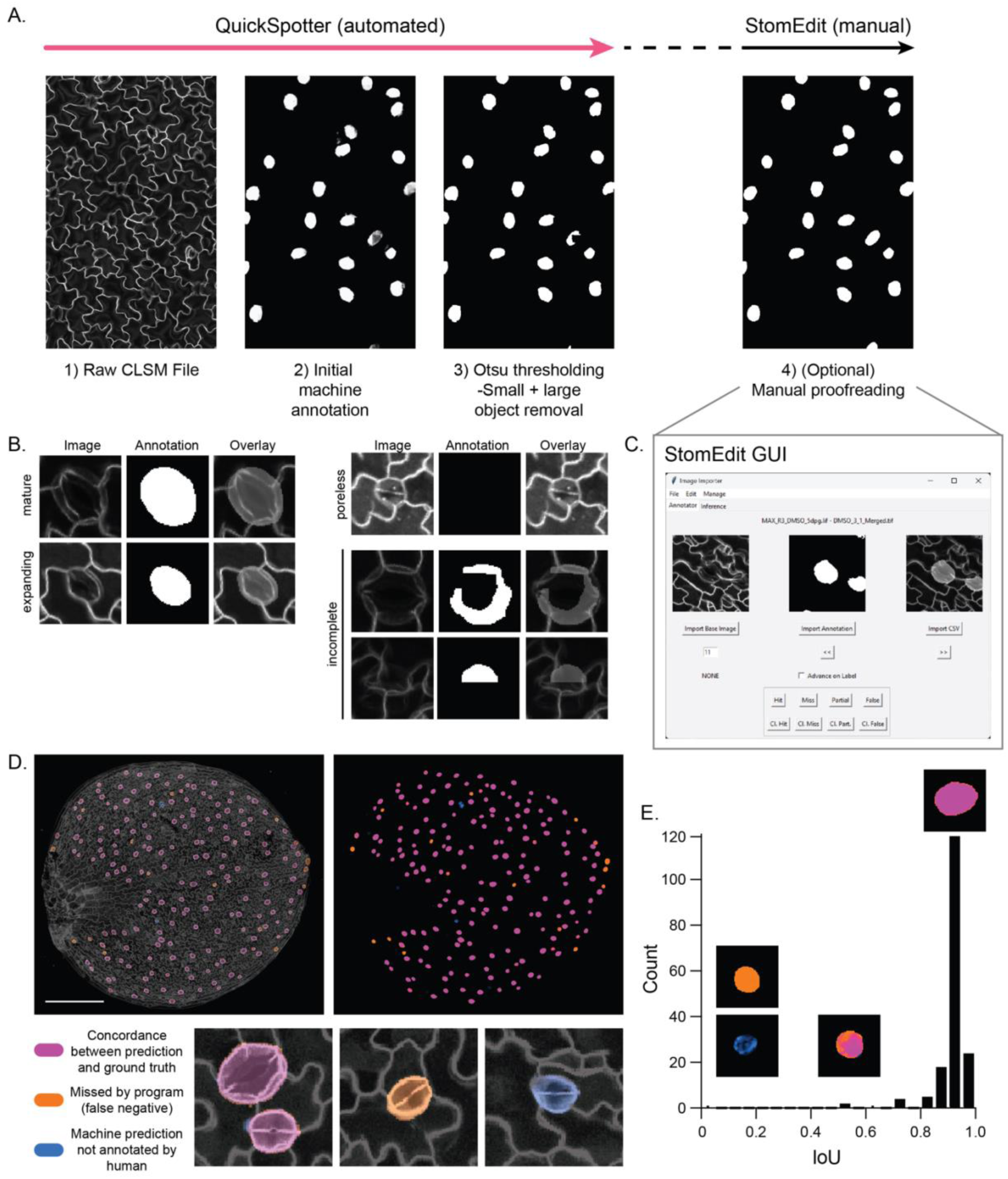
QuickSpotter and StomEdit workflow and associated accuracy metrics. A. Illustration of the QuickSpotter image processing pipeline. A raw CLSM file (1) is examined and used to generate an initial grayscale annotation (2), then processed to yield a fully black-and-white image (3). From there, image defects can be repaired using the built-in StomEdit GUI (see C). B. Representative examples of different annotation categories following the QuickSpotter thresholding step. Stomata with visible pores are annotated with the greatest accuracy. Very young stomata may be missed and masks are occasionally incomplete. C. Screenshot of the StomEdit GUI. The raw and thresholded images are automatically composited in the right viewing window, where each annotated stomata can be examined individually and labeled for phenotyping studies. D. Visual representation of concordance in a 5dpg Col-0 cotyledon between the automatically generated masks and human-generated ground truth. Machine guesses are shown in BLUE, while human annotations on the same leaf are shown in ORANGE. The areas were they concord, pixel for pixel, are shown in pink. E. Bar chart of IoU/contingency errors over all annotations found between the human and machine-generated annotations on a single cotyledon. Representative examples of annotations are shown above their corresponding IoU scores.

**Table 3.** IoU (contingency) statistics for a sample dataset. Three full-size basl-2 mutant cotyledons at 5dpg (B0-B2), five full-size wild-type cotyledons at 3, 4, and 5dpg (C0-C4), and five segments of wild-type cotyledons (S0-S4). The IDs marked with asterisks contain portions of annotations on the edge of the image that were present in the machine guess that were not marked in the ground truth dataset. The “# of margin pixels” value indicates a small region on the boundary of all marked clumps where some inaccuracy is permitted (i.e. a black pixel being annotated as white or vice versa). The “total pixel error” is the number of discrepant pixels between the two images. The “fold error magnitude” is the total pixel error divided by the number of margin pixels, and a fold error magnitude below 1 is considered acceptable.

| Image ID | Contingency | # of annotated pixels | # of margin pixels | Total Pixel Error | Fold Error Magnitude |
| --- | --- | --- | --- | --- | --- |
| B0 | 88.4% | 133939 | 39217 | 15557 | 0.4 |
| B1 | 83.2% | 95465 | 29195 | 16062 | 0.6 |
| B2 | 83.2% | 213280 | 61797 | 35914 | 0.6 |
| C0 | 92.8% | 112785 | 32396 | 8135 | 0.3 |
| C1 | 92.9% | 278316 | 77111 | 19768 | 0.3 |
| C2 | 93.2% | 132810 | 37805 | 9093 | 0.2 |
| C3 | 88.8% | 194832 | 56892 | 21838 | 0.4 |
| C4 | 89.0% | 276002 | 82750 | 30268 | 0.4 |
| S0 | 90.2% | 18678 | 6577 | 1837 | 0.3 |
| S1 | 95.1% | 7395 | 1196 | 360 | 0.3 |
| S2* | 66.4% | 21077 | 6577 | 7087 | 1.1 |
| S3* | 62.4% | 3962 | 1196 | 1489 | 1.2 |
| S4* | 63.5% | 12809 | 4198 | 4669 | 1.1 |

### Graphical user interface for optional proofreading of machine-derived predictions

Even though the accuracy of these machine-generated predictions will likely be sufficient for most applications, we created a tool that could be used to visually examine and correct individual masks if pixel-perfect annotations are required (Figure 2A,C). The StomEdit GUI has several important features for detailed and efficient proofreading of “cleaned” masks, composed of only 0 and 255 values. The GUI gives the user the ability to inspect each stomatal annotation individually, with records that accurately record the position, label and physical characteristics of the annotation. The user can use the GUI to freely mark or erase pixels within each annotation or within an arbitrary region of the image. Deleting a false annotation is expedited within the GUI, as “positive” pixels can be deleted en masse from a given annotation. With these features, using the GUI to proofread a mask of about ∼600 stomata would take only 45-60 minutes in the hands of a skilled user, significantly decreasing the amount of time required to generate pixel-perfect masks for large datasets.

In summary, we have created a machine-learning program that is capable of highly accurate identification and segmentation of individual stomata from fluorescence images. In addition to basic annotations, we have built accompanying tools that can be used to proofread the program predictions and modify masks as needed through a user-friendly GUI.

### Characterizing stomatal changes over cotyledon development

Having developed and validated QuickSpotter, we used it to quantify stomatal characteristics over a critical period of development in 3-7dpg cotyledons. Because stomatal development is asynchronous across the *Arabidopsis* cotyledon epidermis, large sample sizes are needed to accurately capture the range of stomatal lineage progression at each developmental stage. Our program was well-suited to this challenge, capturing the full range of young and mature stomata at each time point. As expected, stomatal counts and density increased from 3 to 7dpg (Figure 3A-C). Average stomatal area and perimeter do not significantly increase with days-post-germination, although the standard deviation of the distribution narrows as a larger fraction of stomata reaches maturity (Figure 3D,E, Figure S2). Interestingly, stomatal eccentricity increased over cotyledon development owing to guard cell elongation during stomatal maturation (Figure 3F-H) (Balcerowicz, 2024; Jaafar *et al*., 2024), highlighting the utility of this program in deriving population-level changes in guard cell morphology from a mixed population of complexes (Supplemental Table 1). Taken together, these data validate the utility of our program and provide a benchmark for population-level stomatal development and maturation in *Arabidopsis* cotyledons.

**Figure 3.**
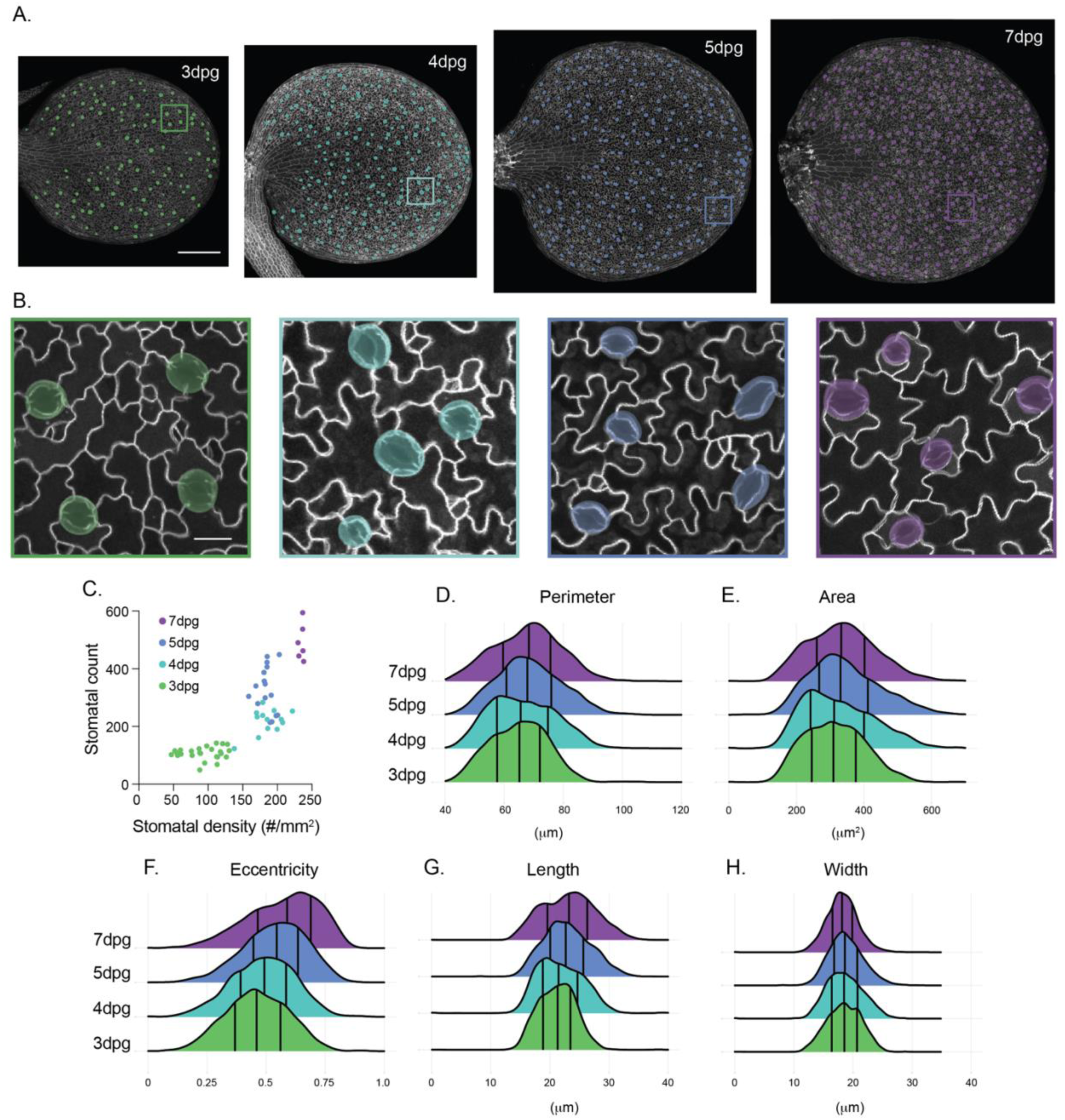
Benchmarking stomatal morphology during cotyledon development using QuickSpotter. A. Representative images of 3, 4, 5 and 7dpg Col-0 cotyledons with associated QuickSpotter-generated masks. Scale bar – 250μm. B. Zoomed regions from each cotyledon in (A). Scale bar – 20μm. C. Scatterplot of stomatal count vs. stomatal surface density calculated from 3, 4, 5 and 7 dpg Col-0 cotyledons. D-H. Ridgeplots of stomatal (D) perimeter, (E) area, (F) eccentricity, (G) length, and (H) width at 3, 4, 5 and 7dpg.

### Changes to stomatal development upon TOR inhibition

Stomatal development is highly flexible and can be tuned by a host of factors, including environmental conditions and nutrient status. One immediate application for our program is systematic analysis of how different perturbations, either through modulation of growth conditions or pharmacological treatment, impact stomatal development and morphology. We decided to apply our program to an image dataset we generated from seedlings that were germinated on plates containing varied concentrations of the Target of Rapamycin (TOR) inhibitor AZD-8055. TOR is a master regulator of growth across plant organs that balances nutrient expenditure with proliferation and differentiation (Caldana *et al*., 2019). Stomatal development is sensitive to sugar availability (Gong *et al*., 2021; Han *et al*., 2022), and TOR-dependent regulation of SPEECHLESS levels has been reported to regulate division potential in the stomatal lineage (Han *et al*., 2022). However, how TOR-mediated regulation of the stomatal lineage is coordinated across developing cotyledons remains unclear.

We grew *Arabidopsis* seedlings on plates containing DMSO or 100nM, 250nM or 1μM AZD-8055, a competitive TOR inhibitor, and imaged their cotyledons at 5dpg. We then used QuickSpotter to analyze roughly 15,000 stomata across the different conditions (Figure 4). We observed a dramatic, dose-dependent effect on overall cotyledon growth, with severe stunting of cotyledon size following treatment with 1μM AZD-8055 (Figure 4A,B). Only treatment with 1μM AZD-8055 significantly affected stomatal counts and stomatal density (Supplemental Table 1). Interestingly, overlaying the stomatal density data following AZD-8055 treatment with those from untreated 3-7dpg Col-0 cotyledons showed that 1μM AZD-8055 treatment caused 5dpg cotyledons to closely resemble 3dpg untreated cotyledons (Figure 4C). AZD-8055 treatment also induced subtle but significant changes in stomatal area and perimeter that made 5dpg stomata more closely resemble stomata from earlier stage cotyledons (Figure 4D-G). Taken together, these data suggest that TOR inhibition slows down overall leaf growth and that division potential in the stomatal lineage is suppressed to keep stomatal density in-line with overall cotyledon area. The reported effect of TOR signaling on SPCH levels may be one avenue to achieve this coordination (Han *et al*., 2022).

**Figure 4.**
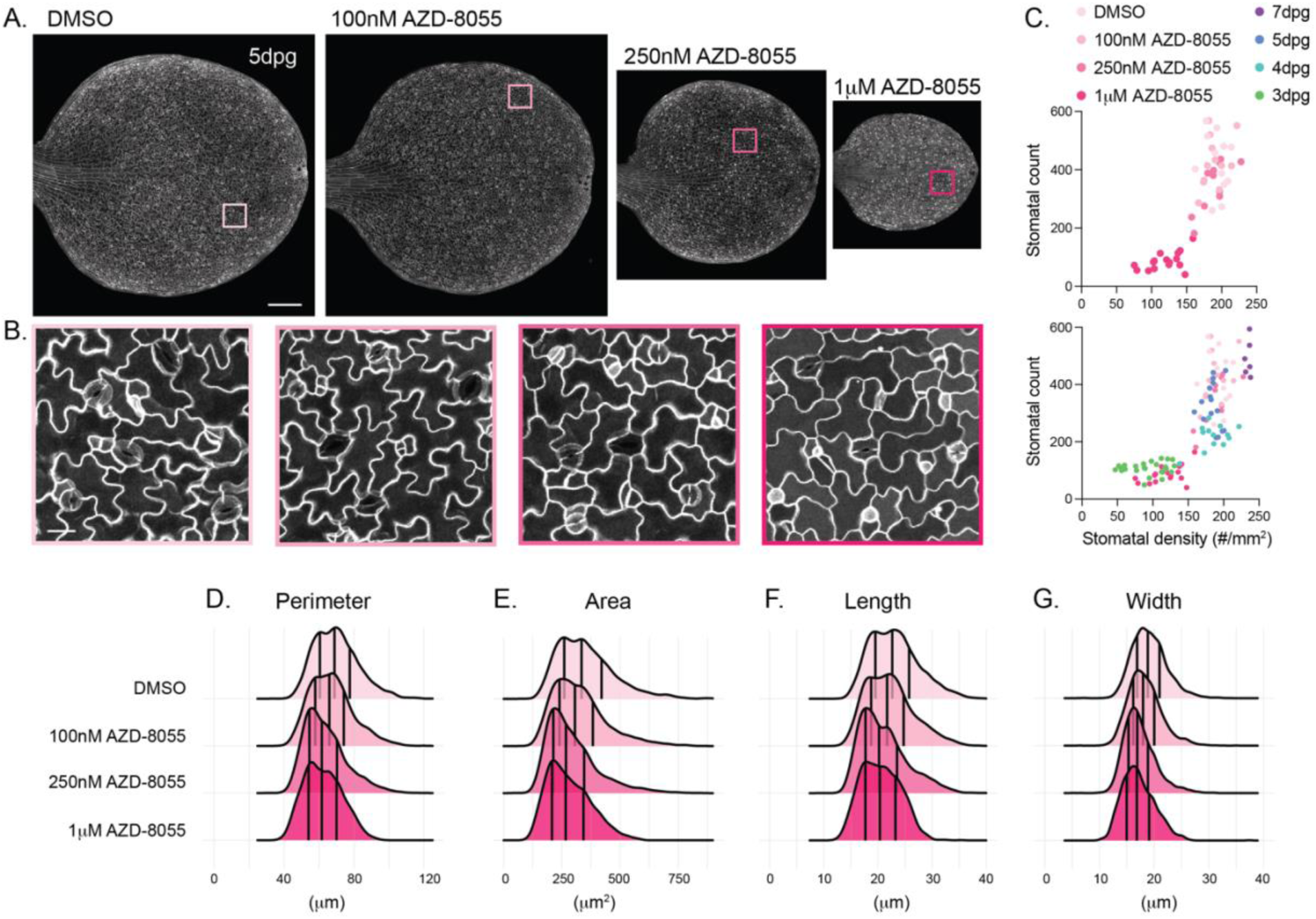
Effects of TOR inhibition on stomatal development and morphology. A. Representative images of 5dpg cotyledons that were grown on plates containing DMSO or increasing concentrations of the TOR inhibitor AZD-8055. Scale bar – 250μm.| B. Zoomed regions on each cotyledon in (A). Scale bar – 20μm. C. Scatterplots of stomatal count vs. stomatal surface density following AZD-8055 treatment alone (top) and alongside data from 3, 4, 5 and 7dpg untreated Col-0 cotyledons (bottom). Note that the data for 3-7dpg cotyledons is the same as those displayed in Figure 3C. D-G. Ridgeplots of stomatal (D) perimeter, (E) area, (F) length and (G) width in DMSO and AZD-8055-treated seedlings.

**Figure 4.**
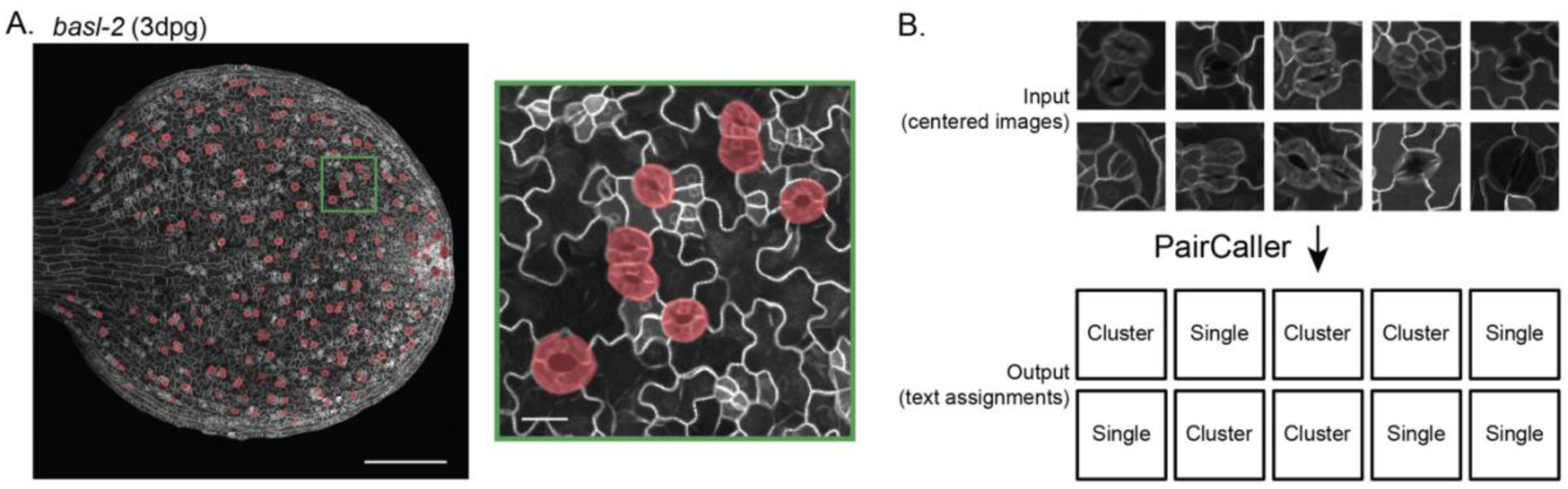
PairCaller accurately categorizes stomata as single or clustered. A. Representative image of a *basl-2* 3dpg cotyledon with QuickSpotter-generated annotations in a red overlay. Note that QuickSpotter does not make a distinction between the annotations that are single stoma versus paired stomata. B. An illustration of the optional PairCaller input and output. The classifier takes as input fixed-size, centered images of stomata, and outputs either a “Clustered” or “Single” label.

#### Add-on functionality to analyze stomatal pairing

In the vast majority of angiosperms, stomata are found distributed throughout the epidermis. This pattern is colloquially known as the “one cell-spacing rule,” and key stomatal regulators, such as EPIDERMAL PATTERNING FACTOR 1 (EPF1), EPF2 and TOO MANY MOUTHS (TMM) (Nadeau & Sack, 2002; Hara *et al*., 2007; Hara *et al*., 2009; Hunt & Gray, 2009), were identified based on their stomatal clustering phenotypes. Furthermore, gas exchange measurements and modeling have shown that pairing has negative effects on stomatal physiology (Lehmann & Or, 2015). Therefore, we decided that an optional add-on to QuickSpotter that evaluates stomatal pairing would have great utility within the community and could be used to identify conditions that trigger subtle but significant increases in pairing.

To train a companion classifier to identify stomatal pairs, we used a series of images from a *BREAKING OF ASYMMETRY IN THE STOMATAL LINEAGE* (*BASL*) loss-of-function mutant. BASL is a polarized protein that regulates asymmetric cell division and stomatal fate in the early stomatal lineage, and *basl* cotyledons and leaves display significantly increased stomatal pairing (Dong *et al*., 2009). Annotating stomatal pairs in this dataset allowed us to develop a sufficiently large ground truth dataset for program training, which would not have been possible using Col-0 data alone (Table 4). Furthermore, applying QuickSpotter to *basl-2* data provided a significant stress test for the program, as early lineage cells (meristemoids, young SLGCs, GMCs) are also increased and clustered in *basl-2,* sometimes resembling young guards.

**Table 4.** Samples used for PairCaller training and validation. Each cell corresponds to the number of stomata present in the training set from a segmented or whole image (rows), arranged by developmental age in dpg (columns) and whether they are present in the training or validation set (training/validation). Single and clustered stomata were included from three images of *basl-2* cotyledons, contributing 312 and 205 stomata to the training and validation groups, respectively.

|  | 3dpg | 4dpg | 5dpg | 7dpg |
| --- | --- | --- | --- | --- |
| Segments | 579 / 25 | 226 / - | 250 / - | 431 / 34 |
| Whole Images | 642 / 112 | 442 / - | 1096 / 208 | - / - |

Over the validation dataset, which included 714 single stomata and 165 clusters of two or more stomata, two clusters were mislabeled as single stomata and 1 single stoma was mislabeled as a cluster. The challenge for a program with this task is to ensure that the classifier examines characteristics of the image intrinsic to single or clustered stomata, rather than finding proxy factors, such as distance of a positively identified pixel close to the image boundary. To limit erroneous classifications of large or single, off-center stoma as clustered stomata, all input images were jittered by shifting them by a random offset. The output of this program is a table of both single and clustered stomata specific to the image under analysis, an associated identification number, and whether the object was classified as a single stoma or a cluster of stomata. These results are elaborated in Table 4.

## Discussion

As essential regulators of photosynthesis and transpiration, stomata continue to be a major research focus across fields of plant biology, including development and physiology. The program we report here will unlock new biological insight by expediting the time-consuming process of stomatal phenotyping. Our machine-vision approach yielded accurate predictions of stomata from fluorescence images, with the optional ability to edit predictions on a pixel-by-pixel basis through a dedicated and accessible graphical user interface. By combining a semantic segmentation approach with the ability to edit files quickly and directly in a purpose-built interface, QuickSpotter can generate precise annotations while also providing the flexibility to fine-tune and diagnose imaging conditions at the start of an experiment. Furthermore, analysis of imaging data with QuickSpotter is considerably faster than manual quantifications, particularly for large imaging datasets or more mature samples, which can include hundreds or thousands of stomata per leaf. A stomatal map for an entire leaf that is 80-90% pixel perfect can be generated in less than 10 minutes and proofread as necessary, representing a significant advance over other machine-learning programs. Finally, it is worth highlighting that QuickSpotter does not require expensive hardware or access to a supercomputer cluster; we have noticed that QuickSpotter’s performance is almost indistinguishable on a range of different mid-level computers, indicating that access to a high-powered processor and GPU are not required to utilize QuickSpotter.

We demonstrated QuickSpotter’s utility by benchmarking stomatal characteristics from thousands of stomata across *Arabidopsis* cotyledon development, providing useful baseline data for future comparison. Additionally, we showed how QuickSpotter can be used in combination with pharmacological perturbations to rapidly analyze new stomatal phenotypes. Here, we treated developing seedlings with the TOR inhibitor AZD-8055 and found that, surprisingly, TOR inhibition decreased stomatal numbers indirectly by altering overall leaf growth. This suggests that *Arabidopsis* stomatal development is more attuned to meeting specific stomatal densities under a given set of growth conditions, rather than monitoring TOR directly to gauge how many stomata are produced.

In the future, we envision that QuickSpotter can be used to generate large phenotypic datasets to understand how stomatal development is tuned by subtle changes in nutrient status, environmental stressors, or small-molecule application. We are also excited by two additional extensions for the program that have been built in but not extensively utilized in the work reported here. First, it is possible to continue improving the network’s capabilities by feeding human-proofed annotations back into the algorithm’s training data, iteratively improving its accuracy in a “bootstrapping” loop. In this way, researchers can develop a copy of QuickSpotter that builds off the extensive built-in training but specifically tailored towards their dataset. Second, future model training could be extended to propidium-iodide-stained images, which would be beneficial for certain experimental setups, for example screening collections of existing mutants, where introgression of a plasma-membrane reporter may be impractical.

Finally, we believe that QuickSpotter’s program architecture is not strictly limited to annotating stomata. With enough training data, QuickSpotter can be configured to detect any feature in a monochrome image, although the quantity of training data required will scale proportionately to the complexity of the target feature. For instance, QuickSpotter may be configured to detect points of intersection between pavement cell walls by annotating white dots over those locations. Therefore, we hope that other researchers will consider QuickSpotter as an entry point to applying machine-learning analyses to their own work, thereby further democratizing this emerging and rapidly evolving technology.

## Acknowledgements

We would like to give special acknowledgement to Kurt Mayer for help with data annotation, Daniel Angres for programming and reproducibility advice, and Kensington Hartman, Bianca Lopez, Lili Follett and other members of the Muroyama lab for advice on experimental design. Funding for this project was provided by a Hellman Fellowship and NIGMS R35GM 150466, both to AM.

---

**Figure S1.**
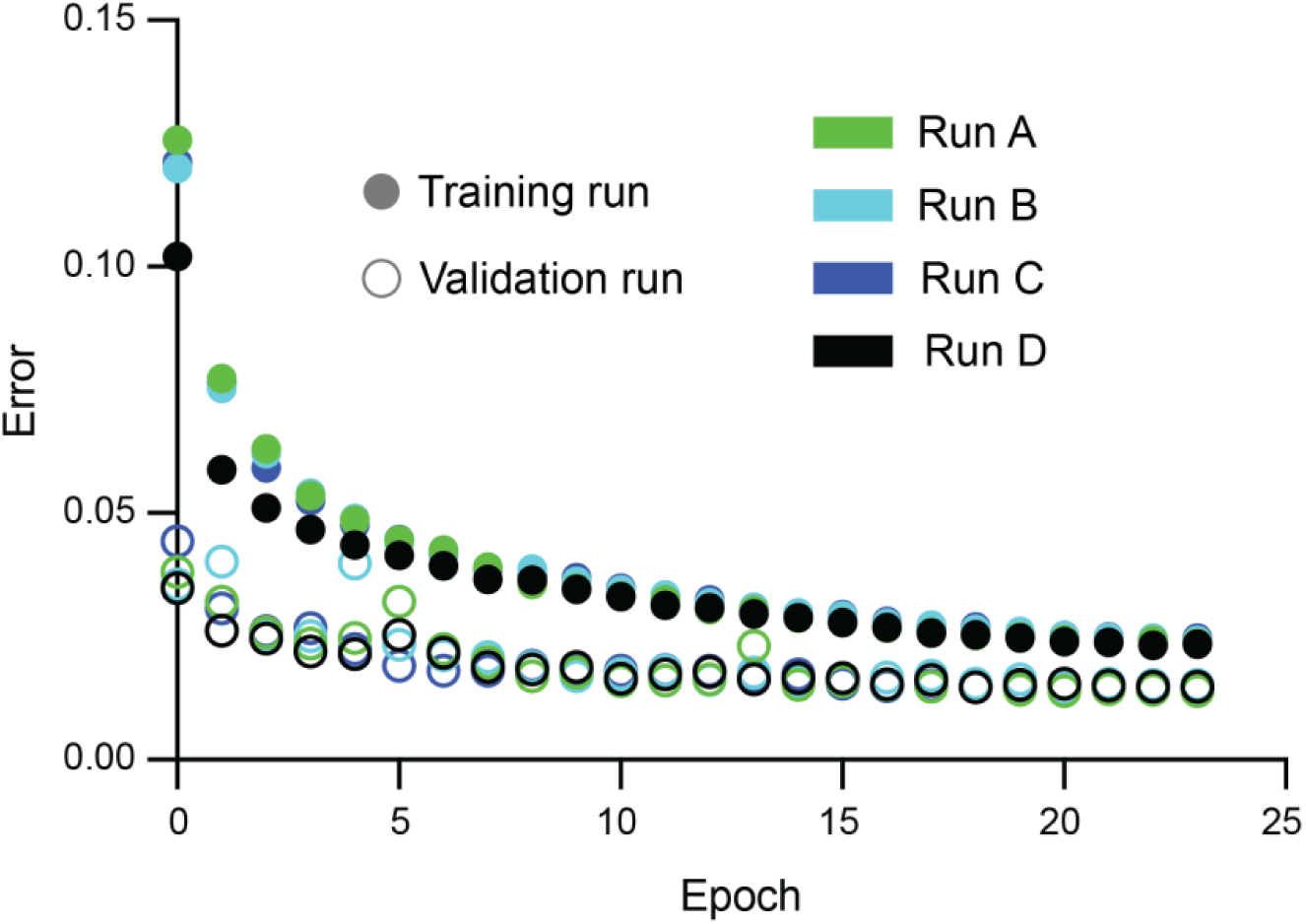
Results from QuickSpotter training cycles. A graph showing a set of training cycles used to verify QuickSpotter’s performance consistency. QuickSpotter’s use of a non-deterministic data augmentation pipeline results in variable fitting performance, since no two input datasets for each epoch will be identical. Here, over four training/validation runs, divergent performance in the early epochs (0-5) converges to common values later in training (15-24). Training and validation runs sharing the same color occurred concurrently.

**Figure S2.**
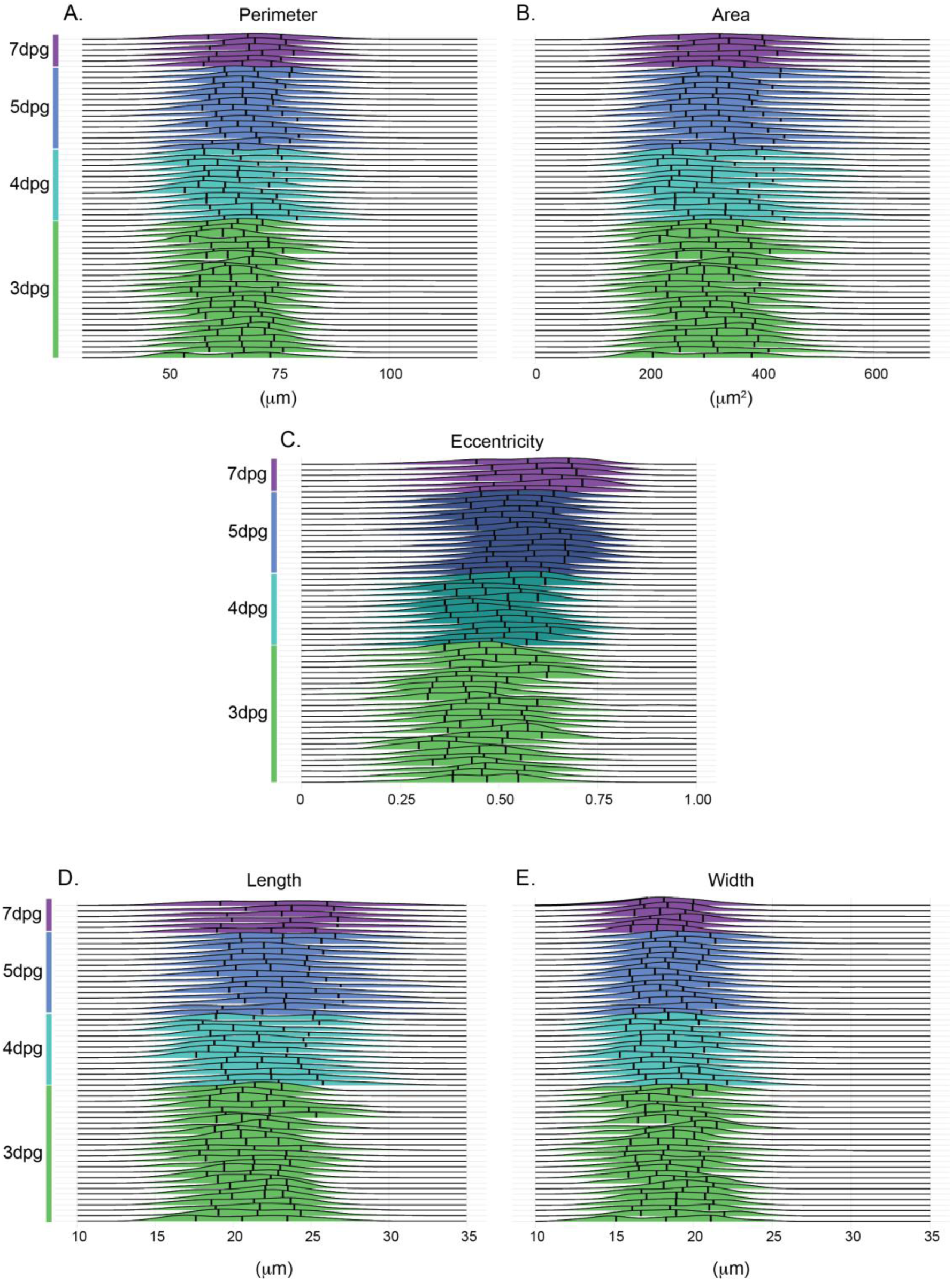
Stomatal morphology across cotyledon development. Ridgeplots of stomatal (A) perimeter, (B) area, (C) eccentricity, (D) length, and (E) width at 3, 4, 5 and 7dpg. Each line shows the stomatal population from a single cotyledon.

**Supplemental Table 1.** Statistics associated with stomatal characteristics across cotyledon development and following TOR inhibition. P-values for the pairwise comparisons between the indicated groups using Kruskal-Wallis tests followed by Dunn’s tests for the post-hoc analysis.

| Comparison | Stomatal count | Stomatal density | Stomatal area | Stomatal perimeter | Eccentricity |
| --- | --- | --- | --- | --- | --- |
| 3 vs. 4dpg | 0.00238 | 0.0000494 | 0.236 | 0.31 | 0.53 |
| 3 vs. 5dpg | 0.000000118 | 0.0000159 | 0.0000104 | 0.00000471 | 0.00000174 |
| 3 vs. 7dpg | 0.000000225 | 0.000000166 | 0.054 | 0.0108 | 0.0000169 |
| 4 vs. 5dpg | 0.603 | 1 | 0.142 | 0.0742 | 0.0236 |
| 4 vs. 7dpg | 0.0538 | 0.255 | 1 | 0.76 | 0.0104 |
| 5 vs. 7dpg | 1 | 0.24 | 1 | 1 | 1 |
| DMSO vs. 100nM AZD | 1 | 1 | 0.419 | 0.446 | 1 |
| DMSO vs. 250nM AZD | 1 | 1 | 0.0000101 | 0.000068 | 1 |
| DMSO vs. 1μM AZD | 0.00000708 | 0.00000382 | 0.00000157 | 0.00000341 | 1 |
| 100nM vs. 250nM AZD | 1 | 1 | 0.0495 | 0.0491 | 1 |
| 100nM vs. 1μM AZD | 0.00000389 | 0.000159 | 0.00794 | 0.0119 | 1 |
| 250nM vs. 1μM AZD | 0.00308 | 0.000228 | 1 | 1 | 1 |

